# DIVA: natural navigation inside 3D images using virtual reality

**DOI:** 10.1101/2020.04.09.019935

**Authors:** Mohamed El Beheiry, Charlotte Godard, Clément Caporal, Valentin Marcon, Cécilia Ostertag, Oumaima Sliti, Sébastien Doutreligne, Stéphane Fournier, Bassam Hajj, Maxime Dahan, Jean-Baptiste Masson

## Abstract

As three-dimensional microscopy becomes commonplace in biological research, there is an increasing need for researchers to be able to view experimental image stacks in a natural three-dimensional viewing context. Through stereoscopy and motion tracking, commercial virtual reality headsets provide a solution to this important visualization challenge by allowing researchers to view volumetric objects in an entirely intuitive fashion. With this motivation, we present DIVA, a user-friendly software tool that automatically creates detailed three-dimensional reconstructions of raw experimental image stacks that are integrated in virtual reality. In DIVA’s immersive virtual environment, users can view, manipulate and perform volumetric measurements on their microscopy images as they would to real physical objects. In contrast to similar solutions, our software provides high-quality volume rendering with native TIFF file compatibility. We benchmark the software with diverse image types including those generated by confocal, light-sheet and electron microscopy. DIVA is available at https://diva.pasteur.fr and will be regularly updated.

## 1. Introduction

Technological advances in the fields of optical and electron microscopy have enhanced our abilities to discern three-dimensional (3D) biological structures via slice-based tomography. Entire organisms can be imaged at sub-cellular resolution and the complex interplay between 3D geometry and biological activity explored [1]. Yet, gaining an intuitive understanding from these complex raw data remains a challenge, as natural modes of 3D visualisation are lacking. Namely, viewing 3D data on a computer monitor while simultaneously using a mouse to interact and extract information is tedious and difficult, *e.g*. clicking inside a 3D object on a 2D screen is a nontrivial task.

Recently, virtual reality (VR) has reemerged as a technology of interest in a host of applications due in large part to new low-cost consumer headsets. Through the seamless integration of stereoscopy, motion tracking and total immersion, VR provides a natural means to visualize 3D structures. Interactions with the aid of VR controllers are intuitive as they are performed as if the data were physically present to the user. Today, numerous initiatives have focused on taking advantage of this technology in the domains of education and scientific research [2, 3, 4, 5, 6]. Recent studies have additionally highlighted the benefits of immersive viewing for handling 3D data which include efficiency and enhanced intuition relative to standard monitor-based visualisation [5, 6]. Various companies and laboratories laboratories have begun to leverage this technology for scientific image visualisation with different volume representation and interaction approaches, a comparison of which is made in Table 1.

**Table 1:** A comparison of currently existing VR solutions.

|  | Arivis In-ViewR | IstoVisio-syGlass | ConfocalVR | VRNT | TeraVR | DIVA |
| --- | --- | --- | --- | --- | --- | --- |
| Native TIFF Compatibility | No | No | No | No | No | Yes |
| System Requirements | High | High | Medium | High | High | Medium |
| Desktop Mode | No | No | No | No | No | Yes |
| Time Series Compatibility | No | No | No | No | No | Yes |
| Applications | Generalized | Generalized | Confocal<br>Mi-<br>croscopy | Neurobiology | Neurobiology | Generalized |
| License | Commercial | Commercial | Free | Free | Free | Free |
| Reference | [7] | [8] | [9] | [10] | [11] |  |

VR is a promising vehicle through which experimentally acquired images can be better understood. It allows instantaneous understanding of complicated 3D morphologies and, notably, through controller-aided interaction it allows entirely natural selection of voxels in 3D space. In our view, image visualisation in VR should be readily available to all research laboratories (as software such as ImageJ and Icy are [12, 13]) and it must provide tools adapted for processing the enormous diversity of microscopy images.

With this motivation, we introduce DIVA, a complete software application that allows easy integration and navigation inside any 3D scientific image using VR. DIVA uses standard 8- and 16-bit TIFF image stacks and hyperstacks (*i.e*. time series images) as input to instantaneously generate interactive volumetric reconstructions. DIVA does not require any image pretreatment nor does it require conversion to intermediate file formats, as is the case in most 3D image visualization software. Accordingly, it can be used to visualise any type of microscopy image regardless of the imaging modality, ranging from confocal to electron microscopy. The following sections introduce the features that distinguish DIVA as a powerful human-data interaction context.

## 2. Implementation

Developed using the Unity game engine (https://www.unity.com/), DIVA includes a dual interface: a desktop mode for viewing on a standard computer monitor and a VR mode where the user enters an entirely artificial immersive environment by means of a VR headset. This design decision stems from the realization that, despite the massive improvement in VR technology, spending large amounts of time in VR space is not a comfortable experience. In DIVA, the VR interface is tailored to efficient scientific analysis of imaging data, while the desktop interface focuses on parameter settings and initial visual screening of the data. Hence, features of the software are implemented in each respective mode separately.

The desktop interface allows the user to modify visualisation parameters such as voxel size and lighting. It also includes a user-friendly transfer function interface (*i.e*. a 3D look-up table), which allows users to specify pixel intensity-dependent opacity and colour by means of absorption and emission curves, respectively, in real-time. In this interface data appears as a 3D volumetric representation on a 2D screen. The volume can be translated, rotated and scaled using the mouse.

In the VR interface, DIVA renders image stacks as virtual “physical objects” or “avatars” in a virtual room environment. The user can grasp the image stack and navigate inside it via physical manipulation using the VR controller (bundled with all commercial VR headsets). A key VR function included in DIVA to address the complexity within biological image volumes is a handheld clipping plane that allows interactive removal of portions of the rendered volume in real-time (see video demonstration in supplementary material). This feature is essential when extracting information from dense images such as the case in electron microscopy. A simple-to-use counting and distance measurement tool is additionally included to allow simple 3D quantification of imaging data. Examples of uses of DIVA for optical and electron microscopy images are shown in Figure 2 and corresponding videos in the supplementary material.

Critical to any VR experience is maintaining a sufficiently high frame rate to avoid delays upon rapid head movement by the user wearing the headset. In practice, this is a significant design constraint, as volume rendering techniques are computationally costly and must be performed twice during each frame in VR (*i.e*. once for each eye). To address this challenges, DIVA renders volumes through a ray-casting approach that can have its sampling resolution easily adjusted in real-time to ensure a sufficient frame rate for the user [15]. Additionally, the resolution of the rendered image is dependent on how far the camera (observer) is; the image is better resolved in regions in which the user is interacting or observing. This has the result of maintaining a high frame rate even on modest computer configurations. Accordingly, DIVA runs comfortably on mid-range laptop computers.

**Figure 1:**
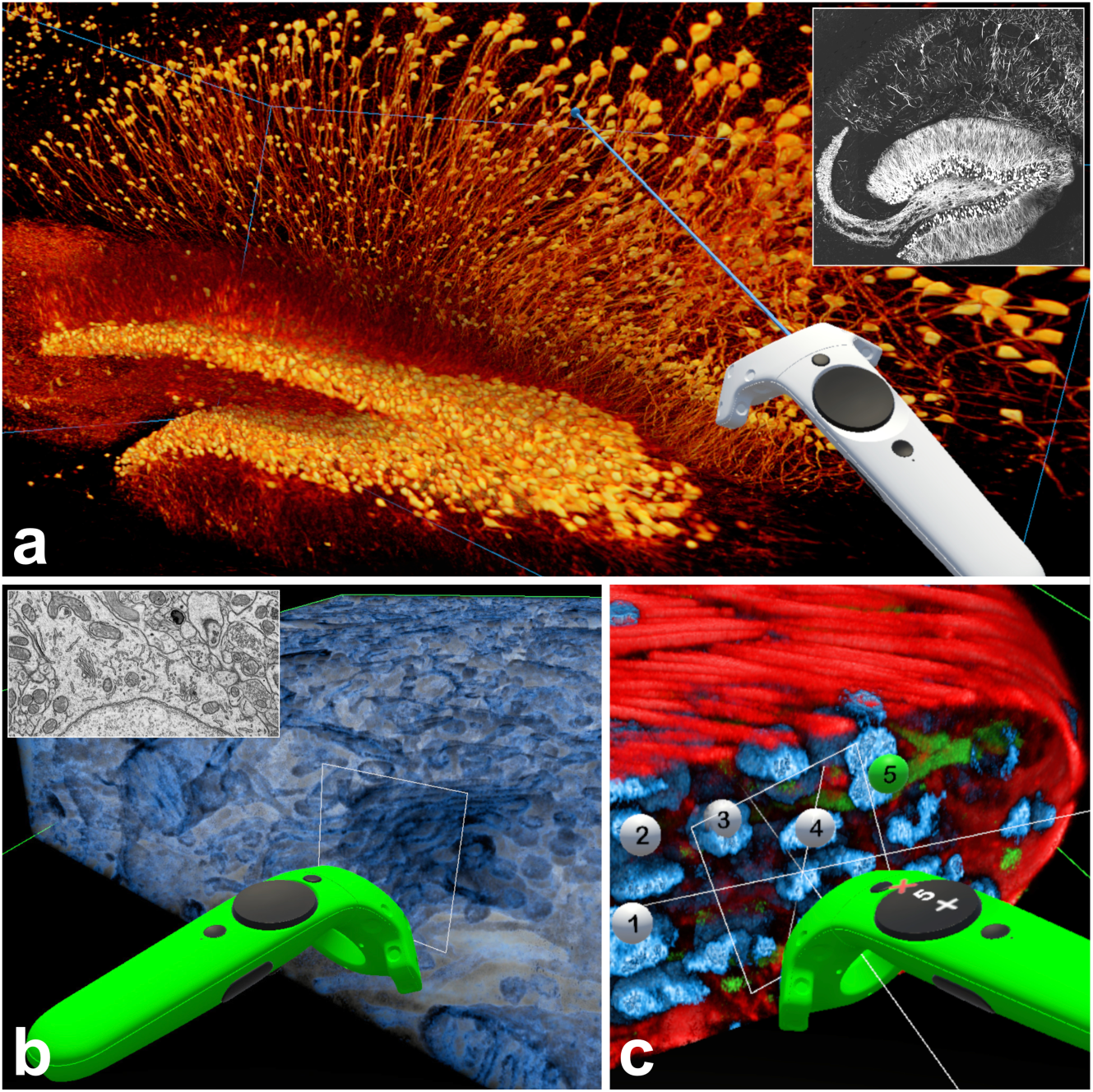
DIVA Usage Examples. Virtual reality visualisation of raw microscopy TIFF image stacks using DIVA. (a) Mouse hippocampus imaged by two-photon confocal microscopy (Thy-1-GFP mouse) with a slice from the raw image shown in inset [14]. (b) Focused ion beam scanning electron microscopy of components of an adult mouse neuron; seen are the Golgi apparatus and mitochondria (raw image is shown in inset). (c) Multichannel confocal acquisition of drosophila testis (red corresponds to actin, blue to nuclei and green to fusome). The positions of nuclei are labeled using a counting tool included with DIVA. In all of the panels, the colour of the VR controller is associated to the action being performed.

Numerous microscopy modalities generate multi-channel or multi-colour image types. Accordingly, DIVA supports visualisation of up to four different channels, whose colour and absorption (transfer function) characteristics can be individually customized. Figure 2(c) shows an example of a multi-colour confocal image rendered in DIVA’s VR context. Additionally, time-series images can be moved where dynamic cellular processes can be observed from every angle using VR. The software has integrated sample images for the user to test upon downloading.

## 3. Applications

To date, DIVA is being used in a number of biological studies. As a pure visualisation tool, it has been used to navigate inside a gamut of microscopy types, not limited to confocal, light-sheet and electron microscopy types. Screen and movie captures are easily made, firmly justifying DIVA’s use as a scientific communication vehicle. Notably, enhanced visualisation with DIVA permits easy quantification and measurements to be performed. For example, filamentous structures such as neurons and microtubules can be readily traced (see supplementary video) and counting of structures in dense images can be done rapidly with the aid of the aforementioned volume clipping plane function. All measurements performed in DIVA can be exported in a CSV format. Table 3 below describes a few of the systems and applications DIVA has been used for to date.

An extensive user manual describing different DIVA use cases is included in the supplementary material. Associated to this are video captures demonstrating the main features of the software.

**Table 2:**
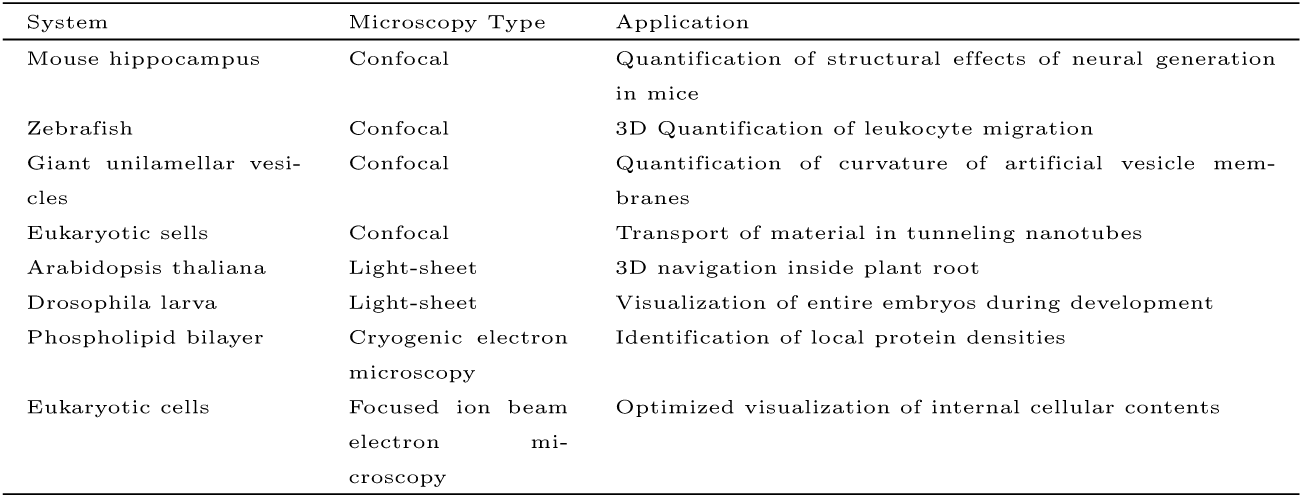
Biological systems that DIVA has been used for to date.

## 4. Conclusion

We have introduced DIVA, a standalone software that allows instantaneous exploration of complex volumetric microscopy images. DIVA leverages virtual reality as a means to explore and “dive in” experimental data in a natural 3D environment. Freely available for academic use, DIVA is a user-friendly software application compatible with all major VR headset brands (e.g. HTC Vive, Oculus Rift) that use the Windows-based SteamVR standard. Updated versions of the software will be made regularly available on https://diva.pasteur.fr.

## Supporting information

Link-to-download-DIVA-viewer

Video Capture of drosophila testes in DIVA

Video Capture of Mice Brain in DIVA

Electron Microscopcy hippocampal cell

caption of the supplementary videos

## Acknowledgments

We thank Kurt Sailor (Institut Pasteur), Richard Weinberg (University of North Carolina) and Thomas Rubin and Jean-Ren Huynh (Collge de France) for the image datasets provided to generate the figures. We acknowledge funding from the Institut Pasteur (J.B.M), The Institut Curie (M.D.), Paris Science Lettres (M.D., J.B.M.) the sponsorship of CRPCEN (J.B.M), the sponsorship of Gilead-Science (J.B.M.), sponsorship of the fondation EDF (J.B.M.), the ANR- 17CE23-0016 TRamWAy (J.B.M),the INCEPTION project (PIA/ANR- 16-CONV-0005, OG) (J.B.M), the “Investissements d’avenir” program under the management of Agence Nationale de la Recherche, reference ANR-19-P3IA-0001 (PRAIRIE 3IA Institute). We thank all the research teams from Institut Curie, Institut Pasteur, College de France who tested the DIVA software and provided feedbacks that have strongly influence the development of the software.

## System Requirements

**DIVA** is designed to run on the **Windows 10** operating system.

Virtual reality (VR) functions require a **SteamVR** compatible headset. Compatibility with such headsets requires, in turn, the installation of **SteamVR** which is done via **<u>Steam</u>**. VR headsets must be calibrated prior to usage with **DIVA**.

To date, **DIVA** has been tested with the following headset models:

- HTC Vive
- HTC Vive Pro
- Oculus Rift
- Windows Mixed Reality (Dell Visor, Lenovo Explorer, etc.)

We recommend using **DIVA** with a modern graphics card with at least 4GB of video memory (VRAM). These may include **NVIDIA GeForce** and **AMD Radeon** models. To ensure stability when running **DIVA**, *it is critical that the latest drivers for the graphics card be installed*.

Note that **DIVA** does not require a VR headset strictly speaking, and can work as a simplified desktop volume viewer.

## Installation

To install **DIVA**, extract the provided zip archive into a desired folder. **DIVA** is executed by double-clicking the provided **diva**.**exe** file.

Important: ***DIVA*** *will take a moment to load as it allocates memory (roughly 20–30 seconds)*.

Depending on installed anti-virus software, notification messages may appear informing of external connections. As an application built using the Unity3D game engine, **DIVA** anonymously sends hardware usage statistics to the Unity server. Although this cannot be disabled within **DIVA**, a workaround is to disable any external internet connections via the **Windows Control Panel** and/or to physically unplug any ethernet cables.

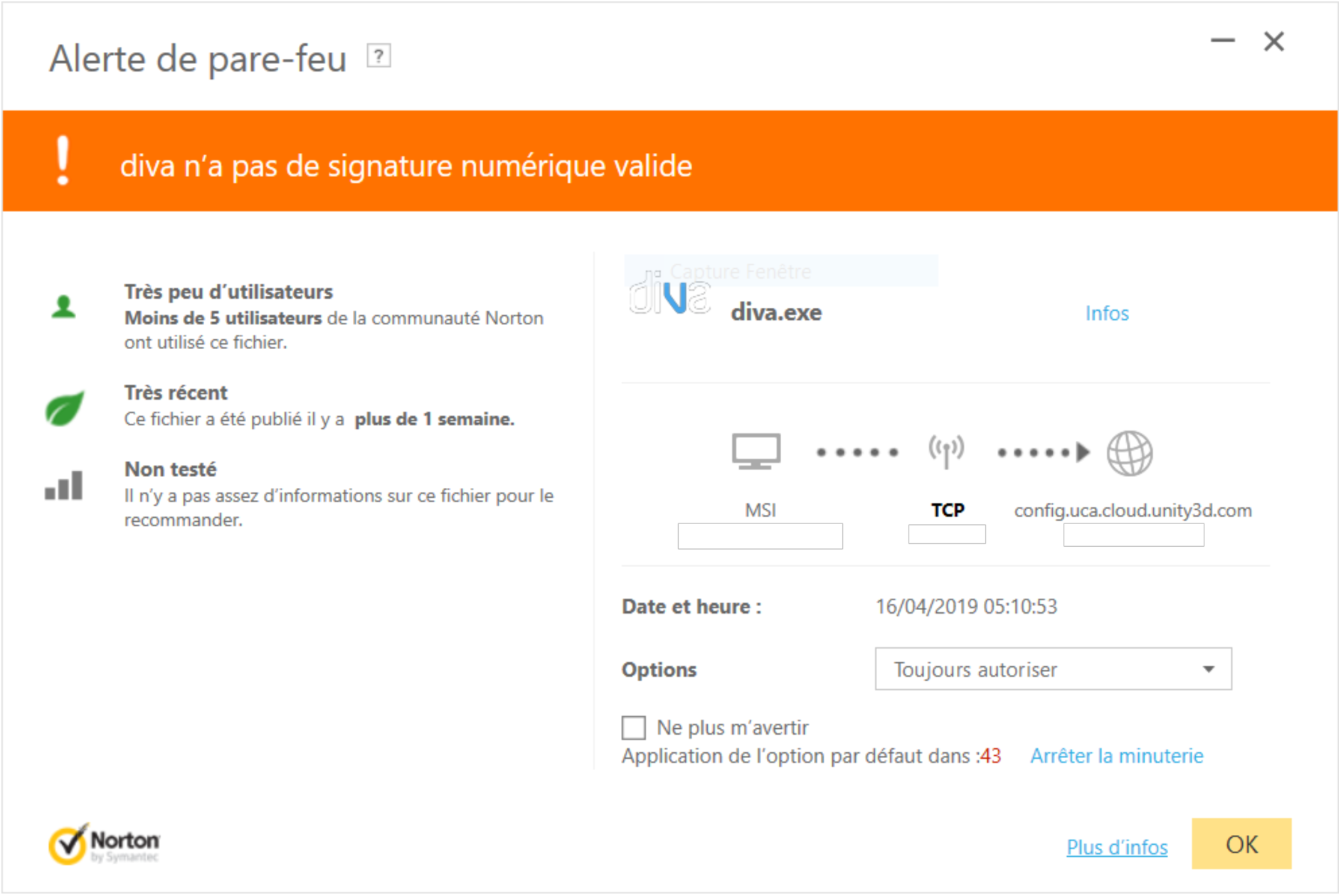

## File Compatibility

**DIVA** processes **Tagged Image File Format** (TIFF) image files. Proprietary scientific image formats (ND2, LSM, etc.) can be rendered compatible with **DIVA** by converting them to a **TIFF** format using for example ImageJ/<u>Fiji</u>.

Multichannel files organized using the ImageJ/Fiji convention are supported. The table below describes which file configurations are specifically compatible.

| Bit Depth | Channel 1 Color | Channel 2 Color | Channel 3 Color | Channel 4 Color |
| --- | --- | --- | --- | --- |
| 8 | Red | Green | Blue | Grey |
| 16 | Red | Green | N/A | N/A |

We recommend limiting the size of loaded files to less than 1 GB. Larger files may be scaled or cropped via ImageJ/Fiji, for example.

**DIVA** is not compatible with movie (e.g. hyperstack) files.

## Desktop Interface

Below is a screen capture of the main **DIVA** desktop interface.

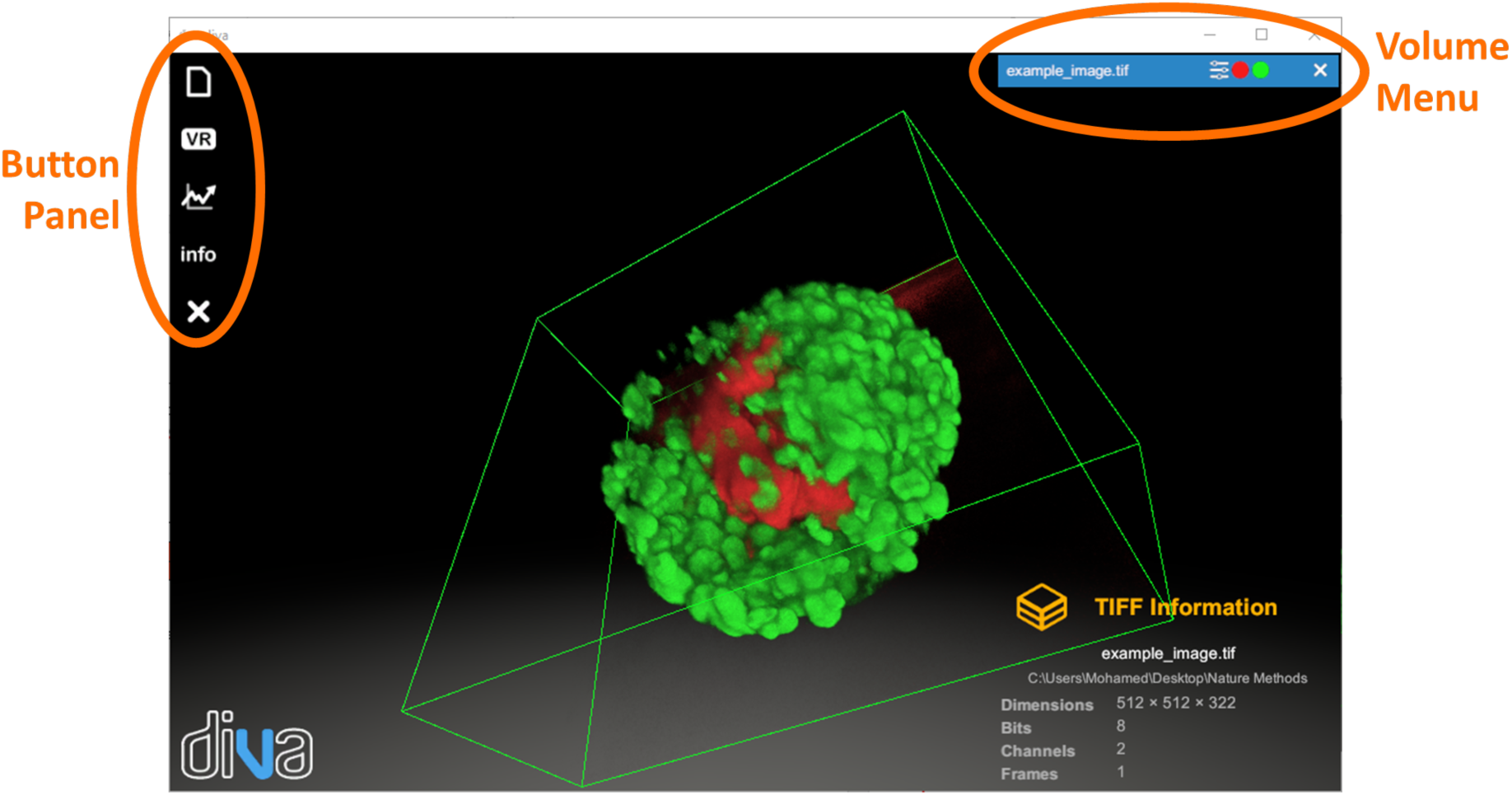

### File Loading

**TIFF** files are loaded by clicking on the button 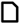 in the button panel. This opens a file browser to select the desired file.

### VR Toggle

Switching to and from VR mode is performed by clicking on the 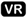 button in the button panel. Clicking this button will automatically launch **SteamVR** to activate the connected VR headset. This button will not respond if **SteamVR** is not installed.

Once VR mode is activated, a feed will be displayed in the VR headset. The desktop will display what is viewed with the headset.

## Results

All screenshots and measurements performed in **DIVA** are summarized in a dynamically generated **HTML Results** file be pressing the 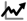 button followed by the 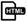 button.

### Visualization Parameters

Visualization parameters for loaded **TIFF** files can be adjusted by clicking on the **Fx** button in the volume menu, a screen capture of which is shown below.

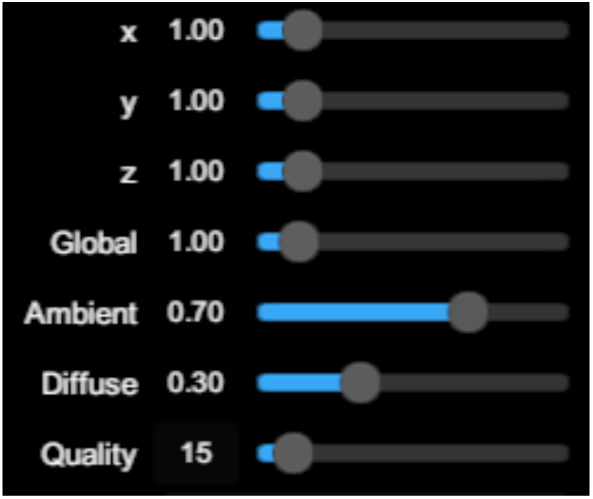

The table below describes the different parameters that can may be adjusted by the user.

| Parameter | Description | Range |
| --- | --- | --- |
| x | Voxel unit size in x dimension | [0.1,10.0] |
| y | Voxel unit size in y dimension | [0.1,10.0] |
| z | Voxel unit size in z dimension | [0.1,10.0] |
| Global | Global isotropic scaling of volume | [0.1,5.0] |
| Ambient | Ambient lighting coefficient | [0.0,1.0] |
| Diffuse | Diffuse lighting coefficient | [0.0,1.0] |
| Quality | Controls quality of rendered volume | [0.0,1.0] |

### Transfer Function

The transfer function, the interface of which is shown below, describes how the loaded image is visualized.

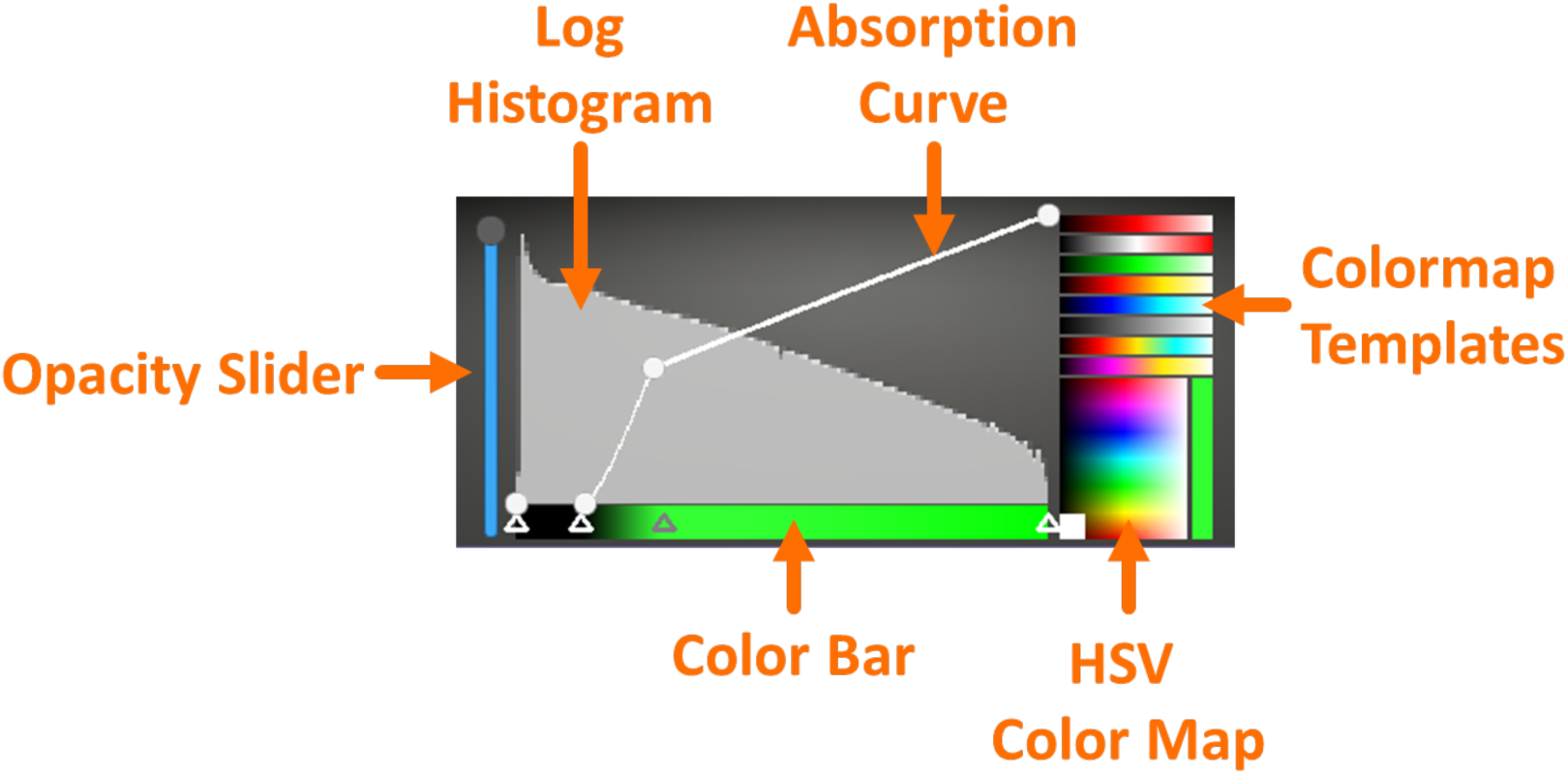

It is composed of a user-defined absorption curve and a color bar, both of which are functions of the voxel intensity (horizontal axis).

The absorption curve defines the opacity of rendered voxels for a given intensity value. It is defined by *control points* which can be adjusted by dragging with the left mouse button. Control points can be added and removed via clicking with the right mouse button. We recommend defining a control point of zero for the lowest intensity pixels as most of the content in loaded images are background (black) pixels.

The color bar allows the user to define colors that may correspond to different voxel intensity values. Similar to the absorption curve, control points are adjusted by dragging with the left mouse button and added or removed by clicking the right mouse button. Colors smoothly vary along a gradient between control points. We recommend using not more than 4 color control points (notably for multichannel images) and setting the lowest intensity pixels to have a black color.

There is an additional slider to the left of the transfer function interface that allows global opacity of the volume to be adjusted.

For multichannel files, each channel possesses its own transfer function which is activated by left clicking on the corresponding channel icon in the volume menu.

Uses of the transfer function interface are included in the provided **DIVA** tutorial videos.

### VR Interface

Once VR mode is activated by pressing the 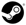 button in the desktop interface, **SteamVR** will load automatically, a screenshot for which is shown below. Note that only one controller and base station is necessary for **DIVA**.

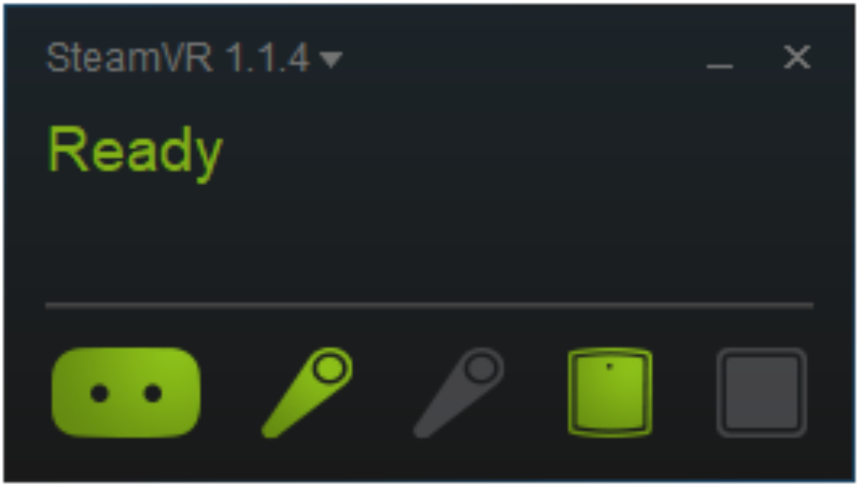

The VR interface is accessed by pointing the VR controller at the loaded volume with the laser pointer and pressing the **Menu Button**, as shown in the figure below. Note that depending on the VR headset being used, the position of the **Menu Button** will differ.

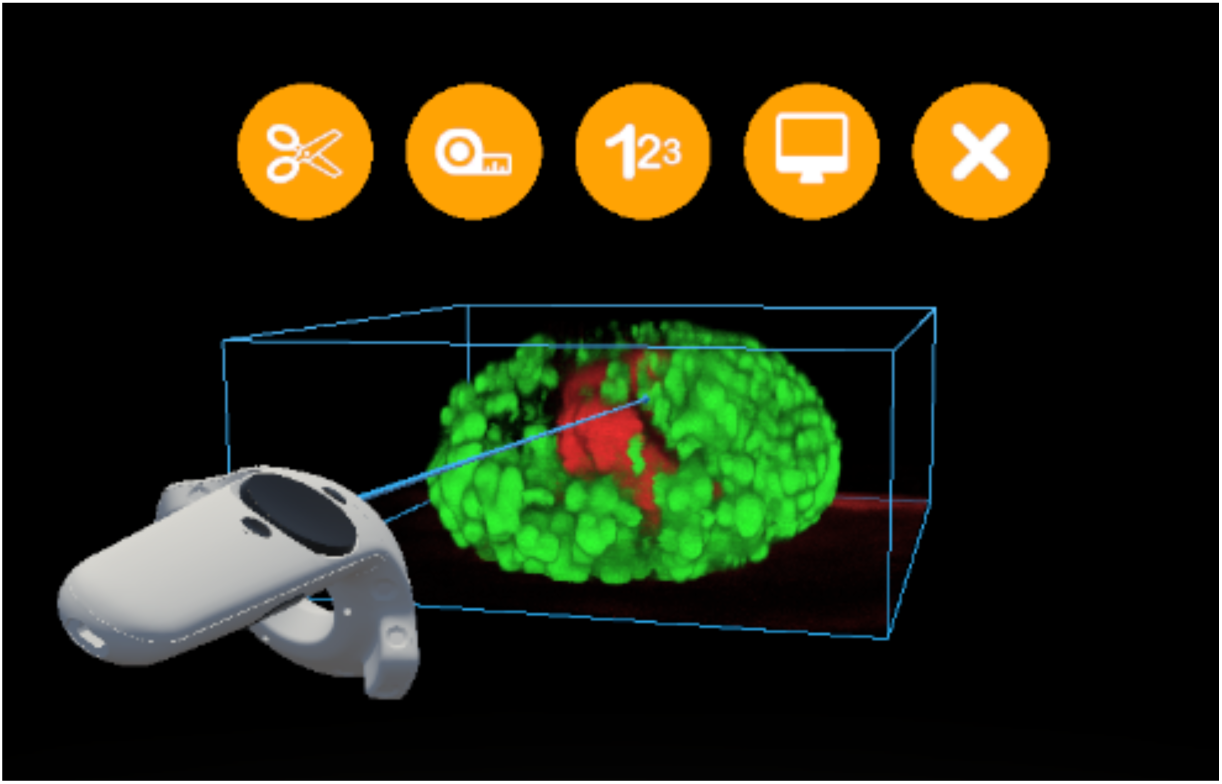

Features of the VR interface menu are accessed by using the VR controller included with the VR headset. Below we display the controller layout for the **HTC Vive** headset. Although initially designed for the **HTC Vive, DIVA** is additionally compatible with other VR headsets (Oculus Rift, Windows Mixed Reality) and button configurations will change accordingly.

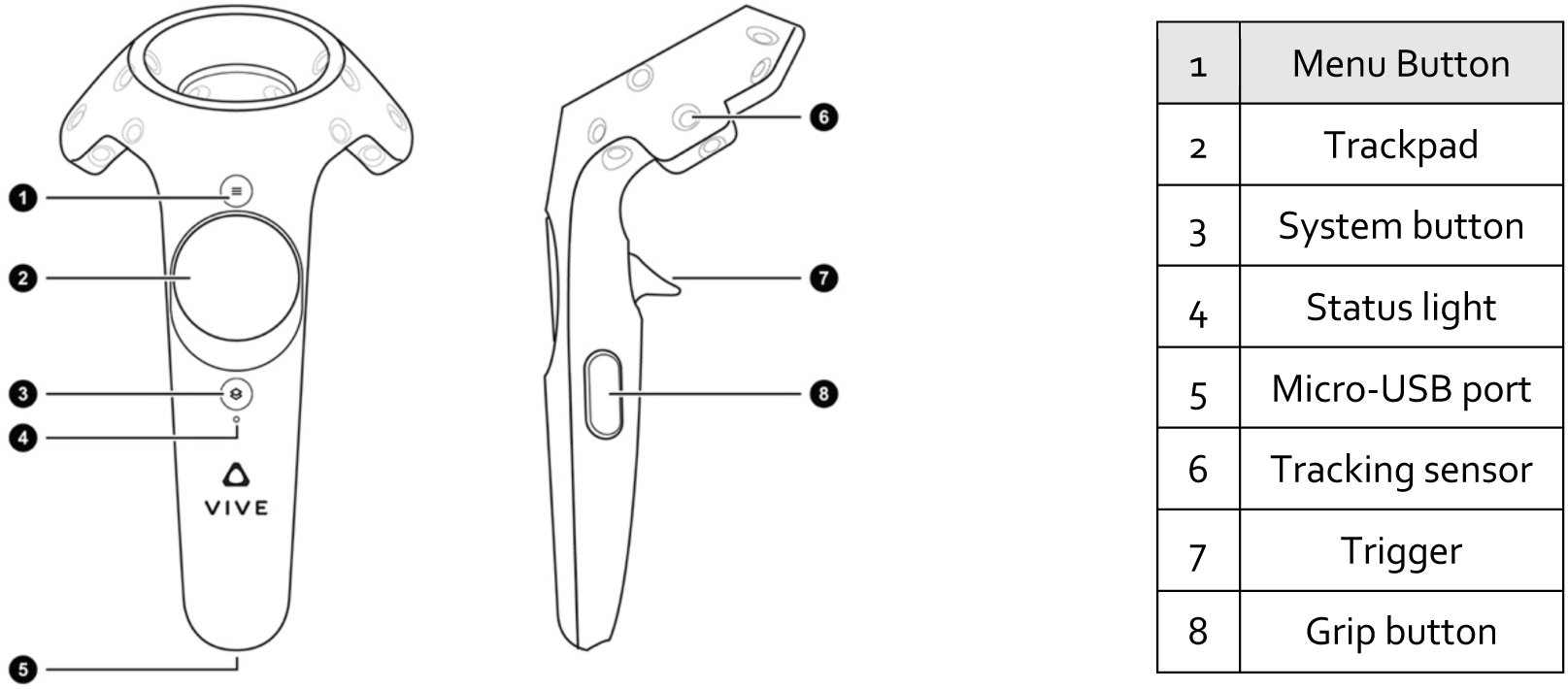

### Physical Manipulation

Once in VR mode, the loaded volume should appear in front of the user. If the volume is not visible, a guide arrow points towards the volume. At any moment, the volume will reappear in front of the user by pressing on the spacebar of the keyboard.

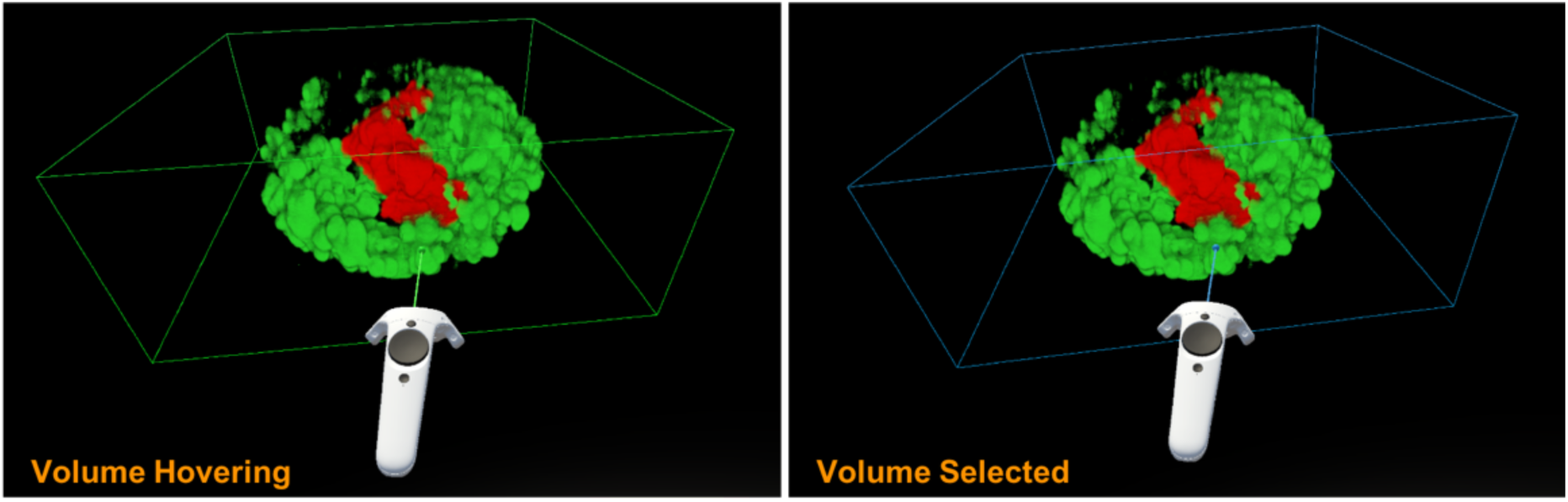

Grasping the loaded volume is done by means of the laser pointer which emanates from the VR controller. The user points at the volume and pressing the trigger button, the volume becomes “attached” to the controller. It can then be reeled, turned and oriented in any direction and position. The volume acts like a physical object; if the user extends their arm with the volume selected, the volume will move away accordingly.

### Clipper

The clipper is toggled by pointing and clicking on the 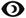 button in the **VR Menu** with the trigger of the VR controller, and then pointing and clicking on the 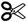 button with the trigger of the VR controller. Upon activation and square appears in front of the VR controller. This feature allows the user to freely clip the loaded 3D volume from any which angle. In order the do this, the user must physically place the controller inside the box surrounding the loaded volume.

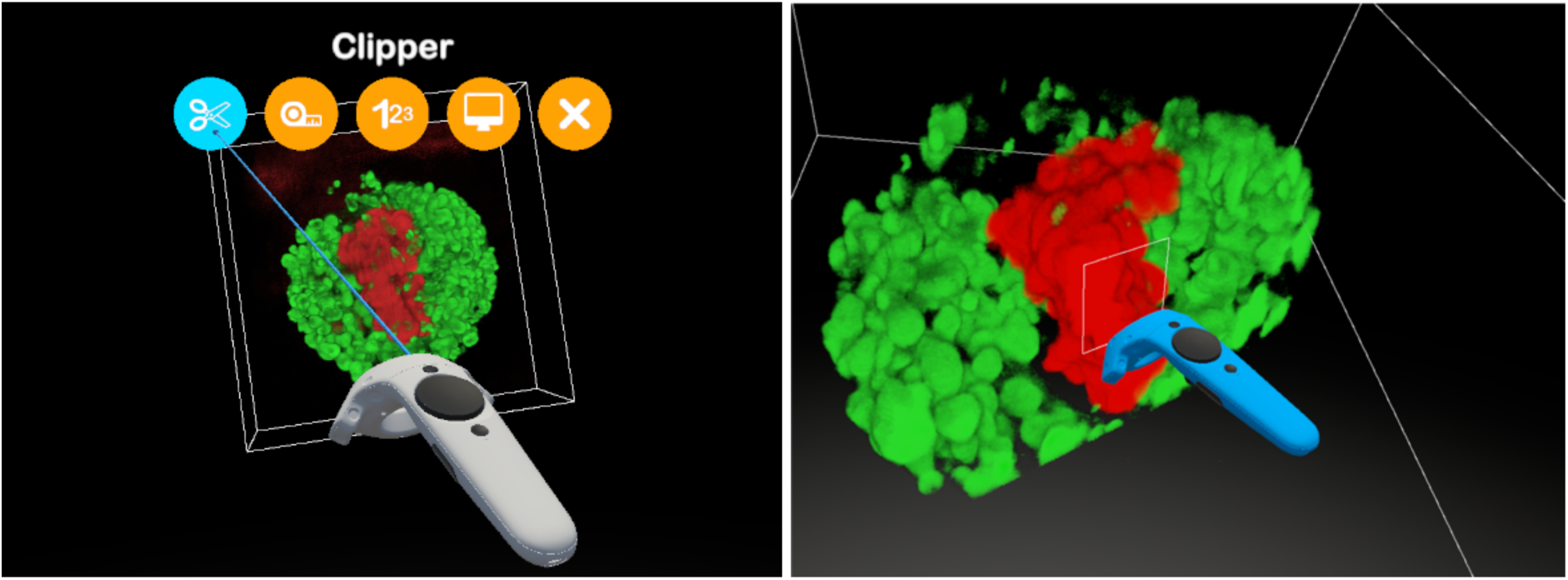

To fix the position of the clipping plane, press any position of the trackpad.

### Ruler

Physical distances within the loaded volume can be measured by means of the **Ruler Interface**. The **Ruler Interface** (below) is toggled by clicking the 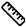 button in the **VR Menu** and pointing at the 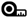 button and clicking the trigger of the VR controller. A specialized widget will appear on the controller that possesses the controls for making measurements, shown below. A small sphere will also appear in front of the controller which designates the position from which distances are calculated.

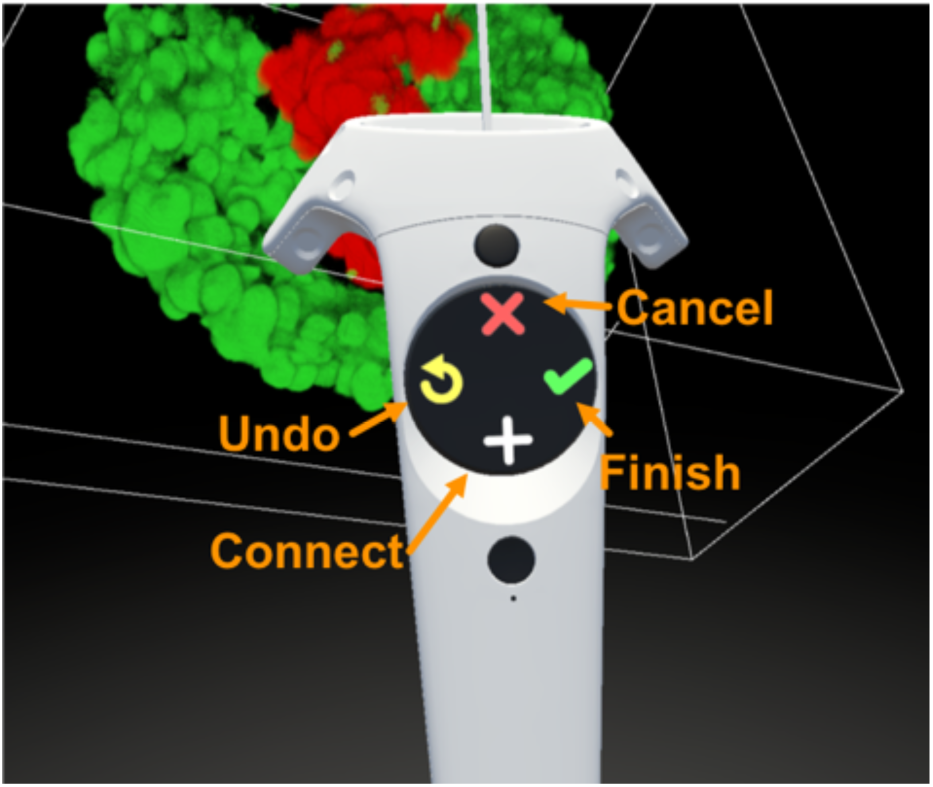

To start a measurement segment, place the small sphere in front of the controller at the desired position and press the **Connect Button** (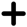 icon). A white line will connect the sphere to the position where the **Connect Button** was clicked.

To connect a segment, simply press the **Connect Button** again. Multiple segments can be connected in this way.

To undo a previously made segment, press the **Undo Button** (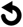 icon).

A measurement is finished by pressing the **Finish Button** (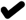 icon).

Finished distance measurements will remain fixed in the volume. The distances of individual segments can be probed by placing the controller in the proximity of a segment until it turns green. Measurement segments can additionally be entirely deleted by pressing the **Cancel Button** (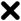 icon) that appears on the controller widget, shown below.

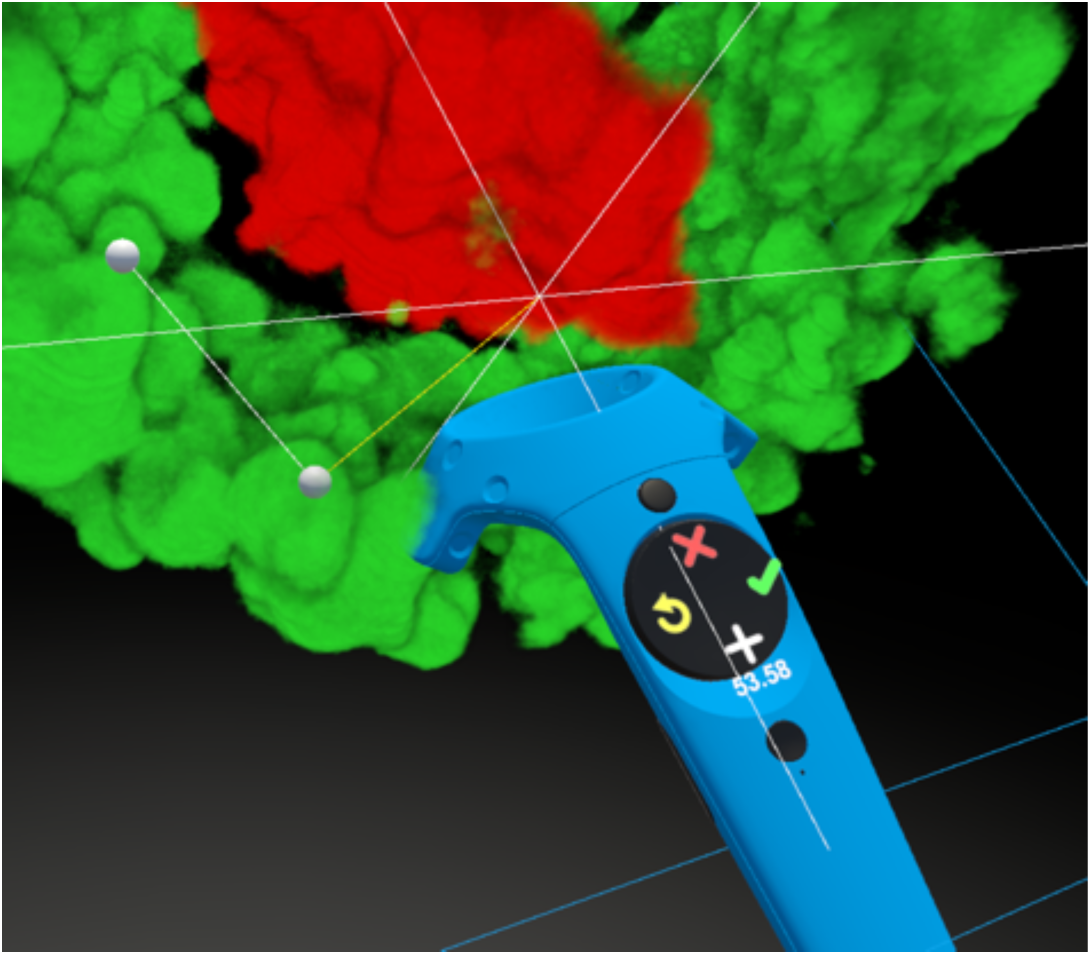

### Counter

Objects inside a loaded volume can be counted by using the **Counter Interface** (below). The **Counter Interface** (below) is toggled by clicking the 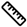 button in the **VR Menu** and pointing at the 1^23^ button and clicking the trigger of the VR controller. A specialized widget will appear on the controller that possesses the controls for counting, shown below. A small sphere will also appear in front of the controller which designates the position where the counter will be placed.

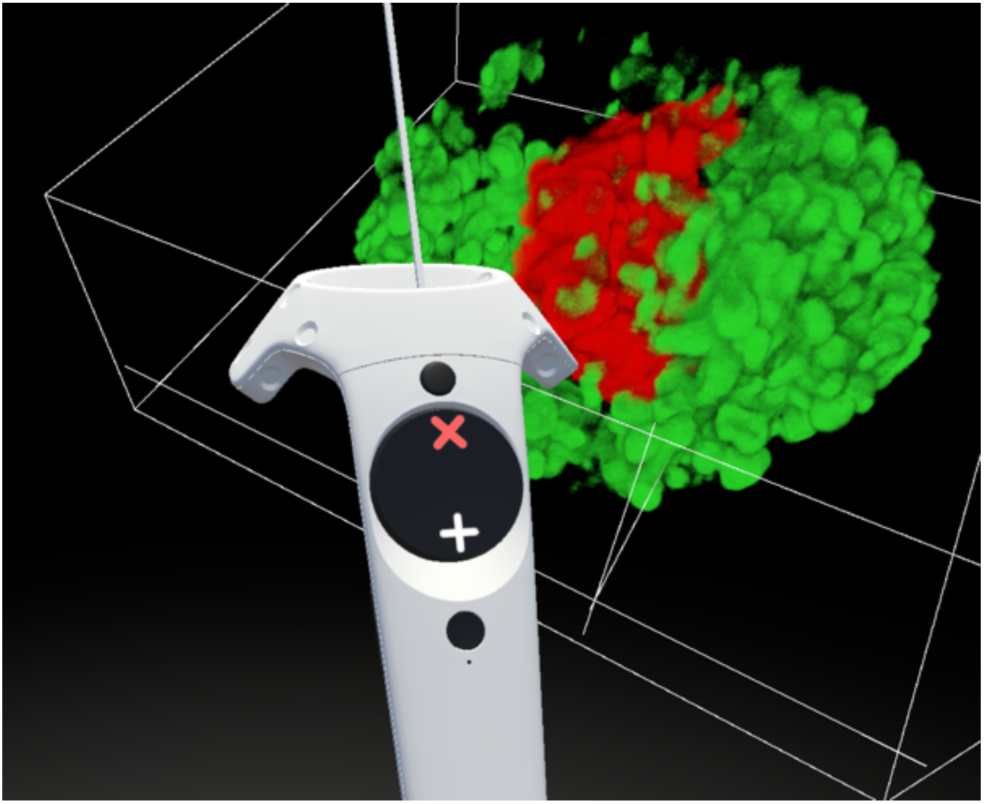

To count, simply place the sphere in front of the VR controller at the desired position, and press the **Count Button** (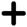 icon). To aid in counting, the **Clipper** can be used in conjunction.

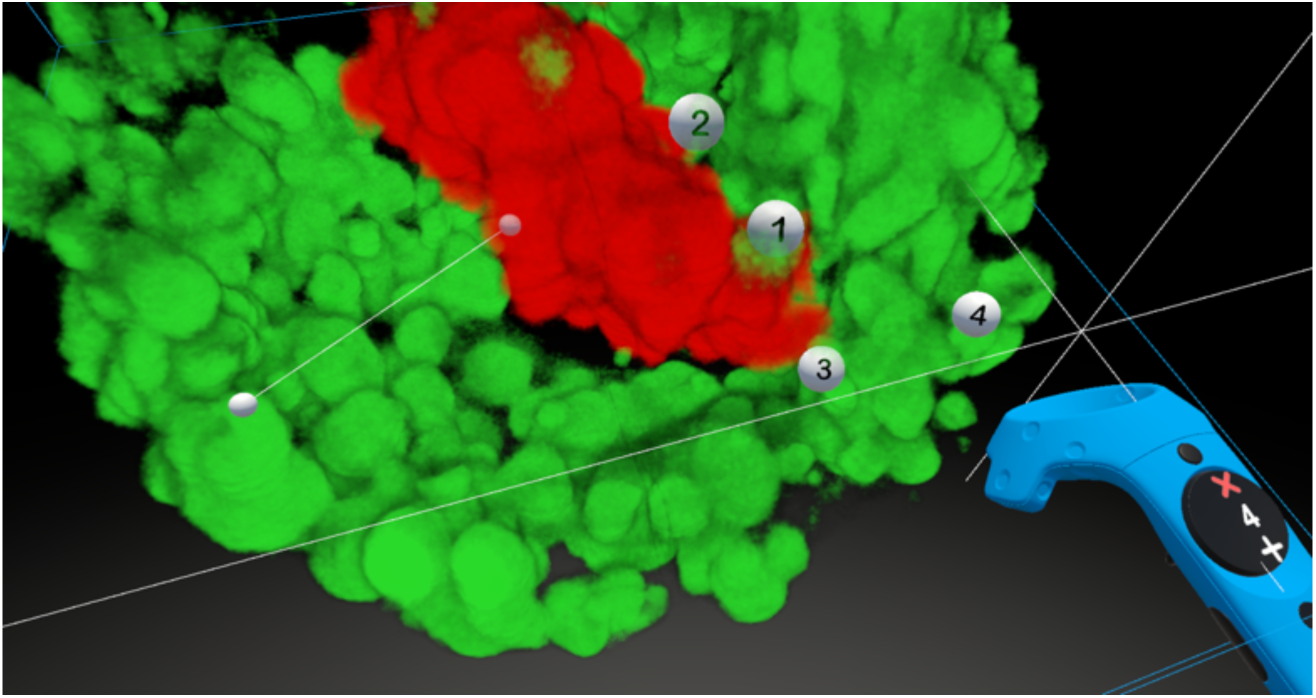

To remove a counter sphere, place the controller in front of the sphere until it changes color to green. At this point, the counter sphere can be deleted by pressing the **Delete Button** (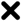 icon).

## Export

### Screen Capture

At any moment when VR mode is active, a screen capture can be made. This is done by pressing the grip button. Screen shots can be retrieved from the **HTML Results** file which is described below.

### Movie Recording

Movies can be recorded at any time when using the DIVA by means of the **Windows Game Bar**. To make a movie recording, hold the **Windows Key + G**. This will load the **Windows Game Bar** interface, shown below, when a screen recording is started and stopped by clicking the red record button.

### CSV Results

Measurements (counts and ruler segments) captured in the VR context can be exported in CSV format. By pressing the 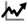 button followed by the 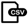 button in the main menu button.

### HTML Results

Measurements and screenshots captured in the VR context are summarized in at HTML file that is dynamically generated by pressing the 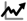 button followed by the 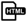 button of the desktop interface. This generates an HTML file that is loaded in the default internet browser, resembling the figure below:

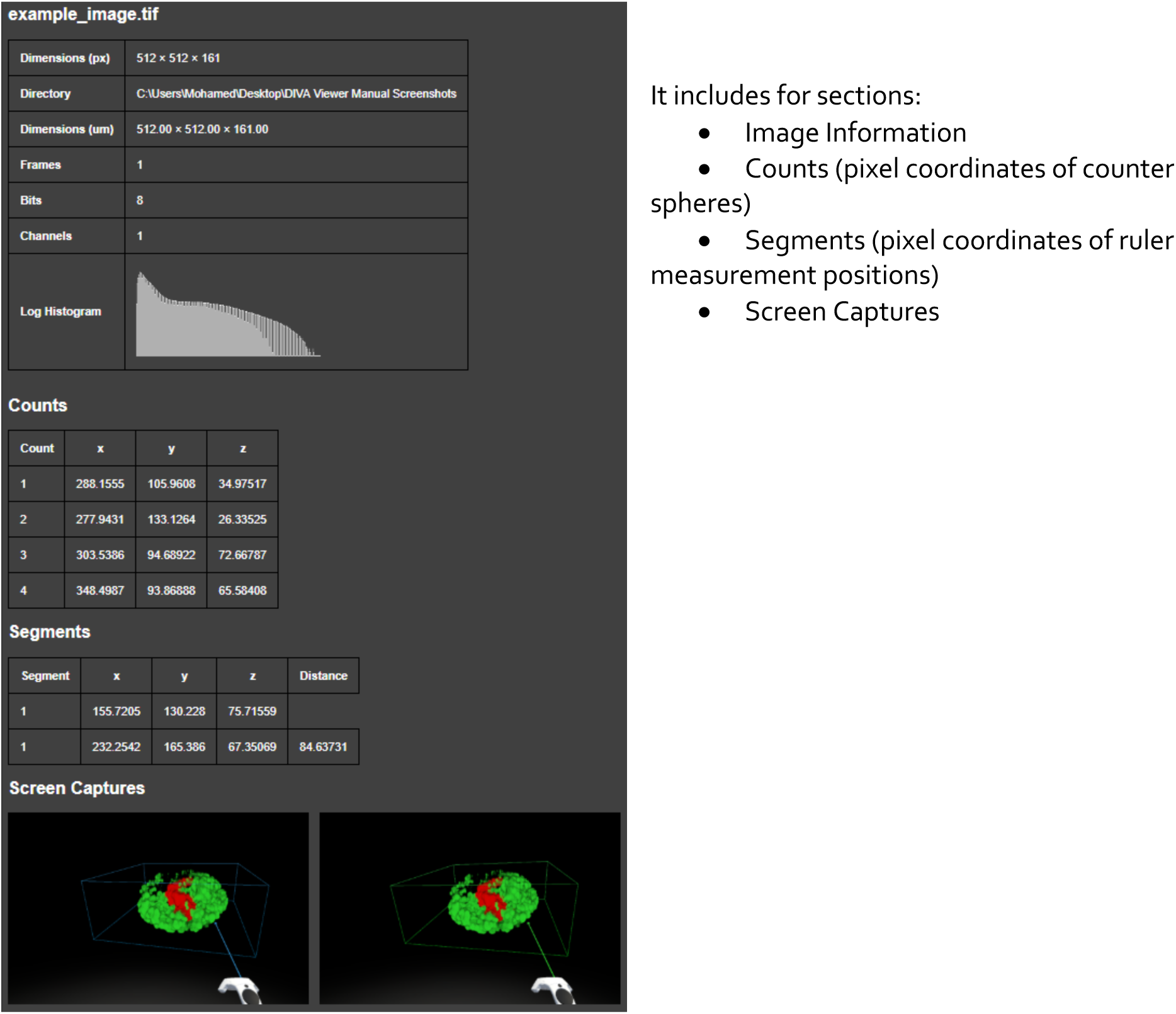

It includes for sections:

- Image Information
- Counts (pixel coordinates of counter spheres)
- Segments (pixel coordinates of ruler measurement positions)
- Screen Captures

## Library Dependencies

External libraries on which DIVA is dependent include the following:

**LibTIFF**

http://www.simplesystems.org/libtiff/

No license

Copyright (c) 1988-1997 Sam Leffler Copyright (c) 1991-1997 Silicon Graphics, Inc.

**SteamVR**

https://www.steamvr.com/en/

**StandAloneFileBrowser**

https://github.com/gkngkc/UnityStandaloneFileBrowser

MIT License

Copyright (c) 2017 Gökhan Gökçe

