## Supplementary material for "DIVA: natural navigation inside 3D images using virtual reality": caption of the supplementary videos

**Video Capture of drosophila testes in DIVA:**

Multichannel confocal acquisition of drosophila testis with the fusomes in green, the germ cell nuclei in blue and the actin envelope in red. Here, DIVA allows seeing structures within structures while accessing the local 3D geometry. VR allows navigation through a tangle of structure with different characteristic geometries, here tubular, spherical and oblong shapes. The interplay of high-density nucleus, fusomes randomly oriented and actin scaffold like structure render the VR visualization necessary to understand 3D spatial organization.

**Video Capture of Mice Brain in DIVA:**

Reconstitution of a serial end block of the whole mice brain imaged in 2-photon microscopy. The tissues are from Thy-1-GFP-M mice that have labeling of subsets of neurons in the brain providing relatively sparse labeling. These mice have enhanced green fluorescent protein under the control of the Thy-1 promoter, which is endogenously expressed in this mouse line. The representation is centered on the anterior/dorsal hippocampus and the densest “v-shaped” structure is the dentate gyrus and the blue dots are the cell bodies of the granule cell neurons. The platform allows visualization in the regions where tagged neurons are sparse with the capacity to follow individual axons along their curved path. It allows also navigating regions with very high density of cell body within the dentate gyrus. Here again, a simple transfer function allowed this visualization without any prior segmenting.

**Video Capture of Hippocampal Neural Electron Microscopy in DIVA:**

Focused ion beam scanning electron microscopy of components of an adult mouse neuron from the hyppocampys. Among the numerous organelles visible, are the Golgi apparatus and mitochondria. Note that there is information in the full volume with no empty voxels. Hence, accessing the full geometry in 3D is challenging as volume perception is hindered by the absence of empty space around organelles. Here, the clipping place allows slicing the volume in real-time and directly detect the mitochondria within the volume. Note the effect induced by the transfer function (look-up table) naturally contouring the organelles without any data preprocessing or segmentation of the volume. DIVA allows characterizing the full structure of the mitochondria in 3D from the EM recording.
